# Overcoming the limitations of motion sensor models by considering dendritic computations

**DOI:** 10.1101/2024.09.09.612048

**Authors:** Raúl Luna, Ignacio Serrano-Pedraza, Marcelo Bertalmío

**Affiliations:** Department of Psychobiology and Methodology for Behavioural Sciences. Faculty of Psychology. Universidad Complutense de Madrid, Madrid, Spain; Institute of Optics. Spanish National Research Council (CSIC), Madrid, Spain; Department of Experimental Psychology. Faculty of Psychology. Universidad Complutense de Madrid, Madrid, Spain

**Keywords:** motion sensors, motion perception, dendritic computations

## Abstract

The estimation of motion is a fundamental process for any sighted animal. Computational models for motion sensors have a long and successful history but they still suffer from fundamental shortcomings, as they disagree with physiological evidence and each model is dedicated to a specific type of motion, which is controversial from a biological standpoint. In this work we propose a new approach for modeling motion sensors that considers dendritic computations, a key aspect for predicting single-neuron responses that had previously been absent from motion models. We show how, by taking into account the dynamic and input-dependent nature of dendritic nonlinearities, our motion sensor model is able to overcome the fundamental limitations of standard approaches.

## 1 Introduction

A major function of the visual system, for any sighted animal, consists of estimating the motion of the elements present in the individual’s environment. This is a fundamental process that guides behaviour, assists navigation, and provides cues that can be essential for survival; for instance, motion estimation is key for visual tasks such as detecting moving objects, appraising distances in our surroundings, segregating foreground from background, and driving eye movements [1, 2, 3, 4].

Abundant experimental evidence suggests that motion sensors are orientation selective, i.e. they can only detect motion orthogonal to their preferred orientation. This motion is one-dimensional, so motion analysis in 2D requires combining the signals from several, 1D, motion sensors [5].

There are two dominant approaches for designing computational models for motion sensors: the earlier one is that of the Reichardt detector [6], which consists in computing the correlation between luminance signals coming from two image locations at slightly different times; the second approach is that of the motion Energy Model [7], based on combining linear filters with nonlinear operations so as to obtain selectivity in space and time.

Two types of motion have been studied extensively: first-order motion, which refers to the movement of luminance features, and second-order motion, where the luminance has no net directional Fourier energy and the motion corresponds to the spatio-temporal modulation of contrast; see Fig. 1 for an illustration. Models that are able to detect first-order motion (such as the seminal Reichardt detector and Energy Model approaches, and elaborations based on them) can not detect second-order motion without the addition of some extra elements, like the combination of motion tracking signals [8], while models that are proposed to detect second-order motion are unable to detect first-order motion. In fact, in [9] it was shown that there is no single model that explains both types of motion. And yet, abundant biological and psychophysical evidence suggests that there are no sensors dedicated only for second-order motion [8, 10, 11, 5, 12, 13].

**Figure 1:**
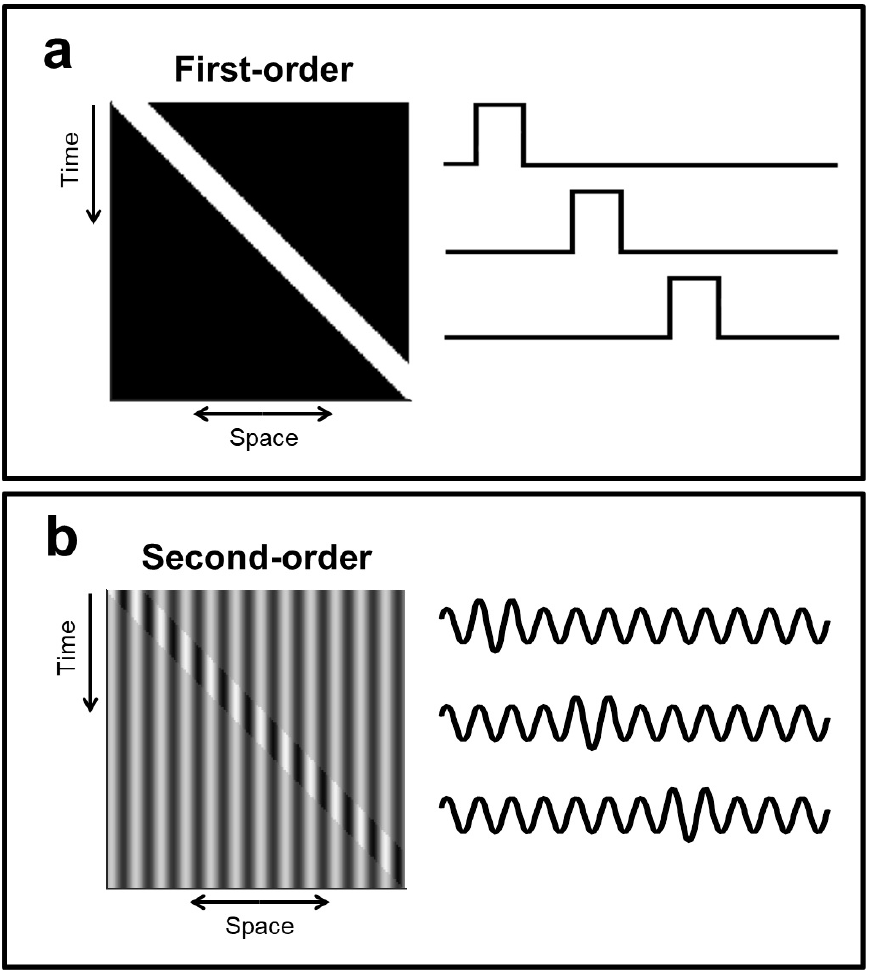
Illustrating the concepts of first-order and second-order motion. **a**, First-order motion, where luminance features change their spatial position. Left: space-time plot of a 1D white bar over black background moving to the right. Right: samples of the 1D signal at different times. **b**, Second-order motion, where the motion corresponds to contrast modulation. Left: space-time plot of a 1D signal consisting of a static sinewave with fixed luminance where a narrow region of increased contrast moves to the right. Right: samples of the 1D signal at different times.

Where are motion sensors located? Motion processing has been observed both in the retina and the cortex of very different species; in the literature, motion sensor models for the retina normally follow the Reichardt approach for insects or the Barlow-Levick model [14] for vertebrates, while the Energy Model approach is favored for the study of motion in the vertebrate visual cortex [15].

Retinal motion detection handles both first and second-order motion, and this is the case both for insects and vertebrates [16, 17, 18], who, perhaps surprisingly, share the same principles for motion sensors at the retina [19]. However, classical motion sensor models do not fit well with retinal physiology [17], have limited accuracy in predicting responses [15] and, very importantly, they do not consider dendritic nonlinearities despite their essential role in providing retinal neurons with direction selectivity, both for vertebrates [20] and insects [21] (of note, direction selective cells have very recently been discovered in the primate retina as well [22]).

In the cortex, both first and second-order motion are processed in the primary visual cortex as well as in extrastriate areas [23, 24, 25, 26, 27, 28, 29, 30]. Experiments have ruled out the Reichardt model as an adequate approach for cortical processing [31]. The Energy Model and all approaches that elaborate on it (e.g. the “filter-rectify-filter” (FRF) model [32], proposed to detect second-order motion and consisting of two phases of energy models with a nonlinear stage between them) are hierarchical, and in the primate they imply a cascaded connectivity going from the lateral geniculate nucleus (LGN) to simple cells in V1, then to complex cells in V1, and then to cells in MT [2]. However, this hierarchical model is not in agreement with physiological evidence. For example, it has been found that MT cells can detect the direction of motion without V1 input [33], and that there is a specialized pathway from LGN to MT, bypassing V1 [34]. Furthermore, it does not seem plausible that neurons of V1, V2, V3, and MT perform either a FRF computation for detecting second-order motion or a linear filtering operation for detecting first-order motion: a more compelling alternative, consistent with principles of cortical processing [35] and with biological evidence [10, 13], is that these neurons located in different cortical areas perform a general and similar type of computation, and that this computation allows them to detect both first-order and second-order motion.

This work proposes a new approach for modeling motion sensors, based on a neural summation model that takes into account the nonlinear, dynamic and input-dependent nature of dendritic computations. The proposed motion sensor model is able to reproduce classical phenomena (e.g. the reverse-phi illusion, motion masking, etc.) while overcoming the fundamental limitations of standard motion sensor models. We will show that the proposed model: (a) is more biologically plausible, as it considers dendritic nonlinearities, a key aspect that had previously been absent from motion sensor models; (b) is compatible with retinal and cortical physiology, and with insect and vertebrate physiology, something that current models lack; (c) is able to detect both first and second-order motion, while current motion sensors detect just one type of motion.

Our results suggest that our proposed motion sensor model is a viable alternative to current models both at retinal and cortical level, as it combines an effective motion detection algorithm with a possible neural circuit implementation that is biologically compelling.

## 2 Results

### 2.1 Proposed motion sensor model

We propose a new type of motion sensor model by extending to the temporal domain a single-neuron spatial summation model recently introduced, called Intrinsically Nonlinear Receptive Field (INRF) [36].

The INRF formulation takes into account several key dendritic properties that are absent from classical models of single-neuron responses, including the following: (a) for the same neuron, some dendrites may act linearly while others may act nonlinearly [37, 38]; (b) there is feedback from the neuron soma to the dendrites, and it can modify the dendritic nonlinearity [39]. INRF is a spatial receptive field (RF) model where the neuronal output is defined in this way:

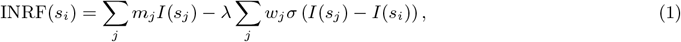

where *s*_*i*_ denotes the 2D coordinates of the neuron location in the image plane and *s*_*j*_ the coordinates of its neighbors, *m, w* are 2D spatial filters, *I* is the static 2D input, *λ* is a weighting parameter (a real number), and *σ*(*·*) is a nonlinear function corresponding to the dendritic nonlinearities. The INRF model was shown to be able to explain very puzzling data in visual neuroscience [40] and visual perception experiments [41], data that challenge the so-called “standard model” of vision that is at the root of motion sensor models, where neuron activity is modeled as a linear summation followed by a nonlinearity [42]; and in artificial neural networks, replacing linear filters with INRF modules results in networks that are more accurate, robust and need just a fraction of the training data [36].

Our proposed extension of the (spatial) INRF model is a spatio-temporal nonlinear RF model that has the following form:

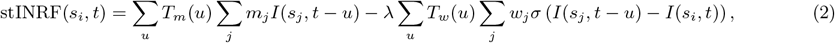

where *t* is the time variable, *T*_*m*_, *T*_*w*_ are 1D temporal filters whose argument *u* denotes a temporal delay, and *I* is now a time-varying 2D input. This stINRF model inherits from the original INRF formulation essential properties that are aligned with retinal and cortical evidence for a variety of species, such as: (a) the inclusion of both passive dendrites with linear responses (in the first term of the model) and active dendrites with nonlinear responses (in the second term); (b) the influence of the input on the dendritic nonlinearities (as the function *σ* is shifted by the local value of the input, *I*(*s*_*i*_, *t*)). But, as a departure from a naïve extension of the INRF model to the temporal domain where the instantaneous response only depended on the instantaneous input, the stINRF model consists in convolving each of the terms of the INRF formulation with a temporal filter: in this way, the stINRF responses depend not only on the current presynaptic inputs but also on the recent activity of those afferents, following [43].

In order to make the stINRF model consistent with physiology as much as possible, we make the following choices, illustrated in Fig. 2:

**Figure 2:**
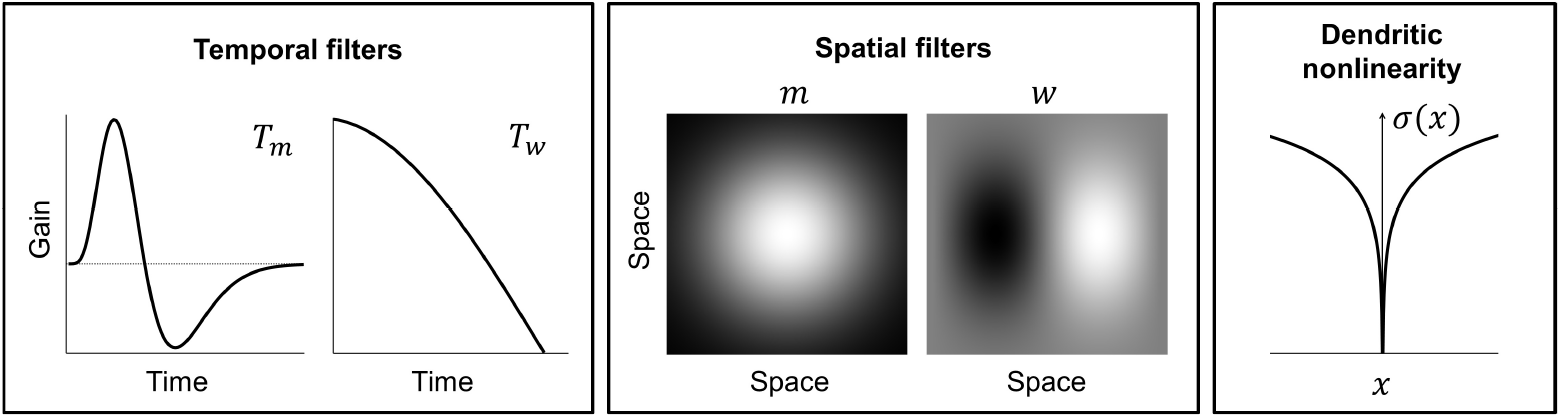
Choices for the elements of the stINRF model. From left to right: bandpass temporal filter *T*_*m*_, lowpass temporal filter *T*_*w*_, Gaussian spatial filter *m*, oriented spatial filter *w* and non-monotonic dendritic nonlinearity *σ*. In order to make the model consistent with biology as much as possible, these choices have been based on a number of works on neurophysiology (see text).

- Based on [44], the temporal filter associated with the linear term, *T*_*m*_, has a bandpass form, while the temporal filter associated with the nonlinear term, *T*_*w*_, has a lowpass form.
- We choose *m* to have a Gaussian shape. And we take *w* to be an oriented filter: aside from the very well-known presence of this type of kernel in the visual cortex, there is evidence of its involvement in performing motion detection in the retina for many different species, from insects to vertebrates, including primates [22].
- Based on very recent results [45, 46], we take the dendritic nonlinearity *σ*(*·*) to be a non-monotonic positive even function.

Finally, the motion sensor (MS) model that we propose consists in computing the mean average (in time) of the stINRF response, i.e. the output of our model for a motion sensor at spatial location *s*_*i*_ and time *t* is computed as the average, over the recent past, of the stINRF output at *s*_*i*_:

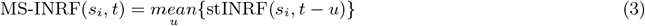

### 2.2 Basic motion sensor properties

The proposed MS-INRF formulation adequately predicts a number of basic visual phenomena that have been reported both for insects and vertebrates:

#### Detection of first-order motion

The MS-INRF model produces motion-opponent responses that are invariant to contrast polarity: model outputs are positive for rightward motion and negative for leftward motion, and these values are insensitive to the polarity of the contrast, i.e. a white bar over a black background and a black bar over a white background will produce the same output if they move in the same way. See Fig. 3.

**Figure 3:**
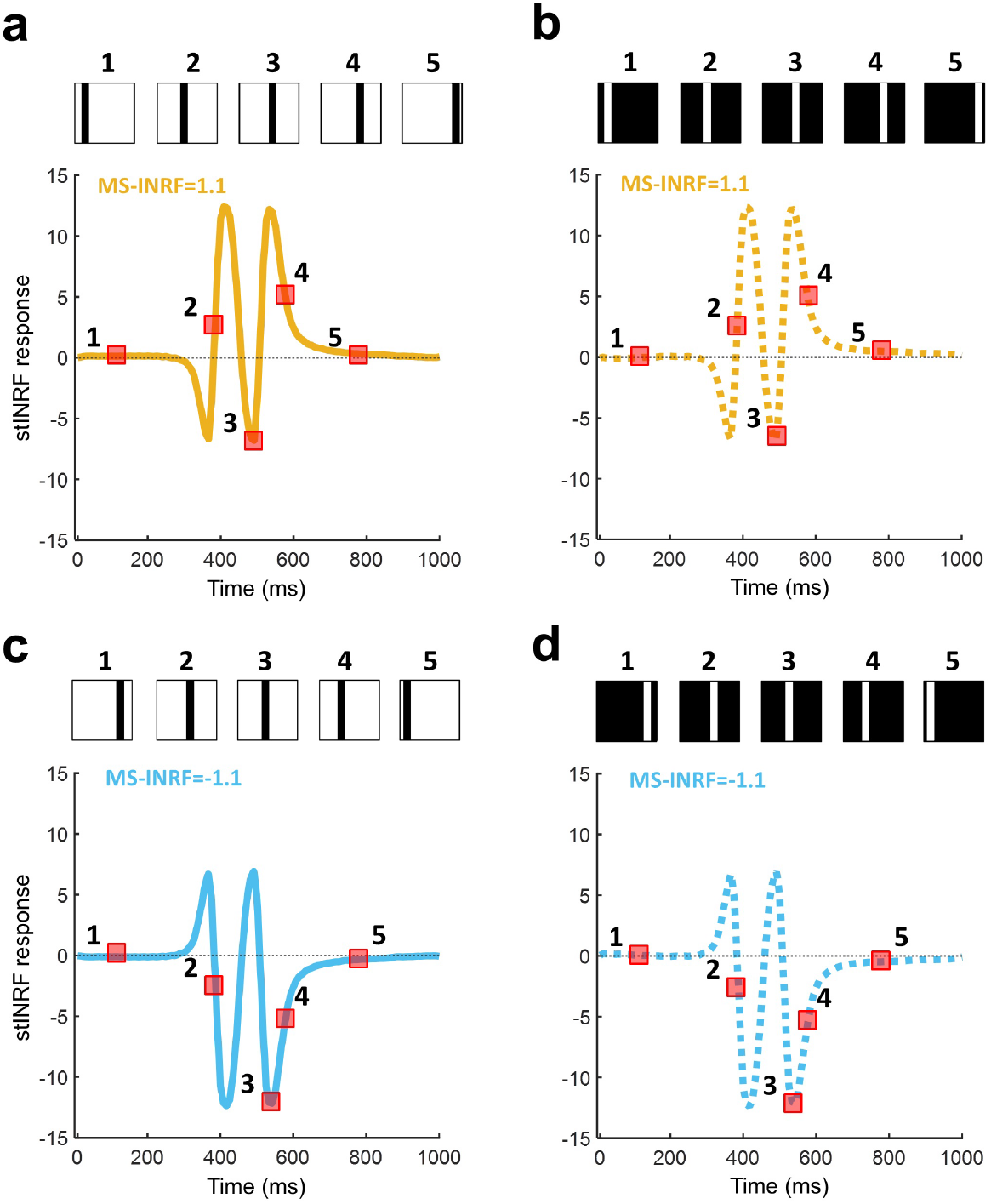
The proposed model MS-INRF produces motion-opponent responses that are invariant to contrast polarity. Motion to the right produces positive MS-INRF responses (panels **a,b**), while motion to the left produces negative MS-INRF responses (panels **c,d**), regardless of whether the moving object is black over a white background (panels **a,c**), or white over a black background (panels **b,d**).

#### Contrast saturation

The magnitude of the model response, as a function of contrast, is monotonically increasing but saturates for high contrast values, see Fig. 4. This is in agreement with classical findings, e.g. [47, 48, 49].

**Figure 4:**
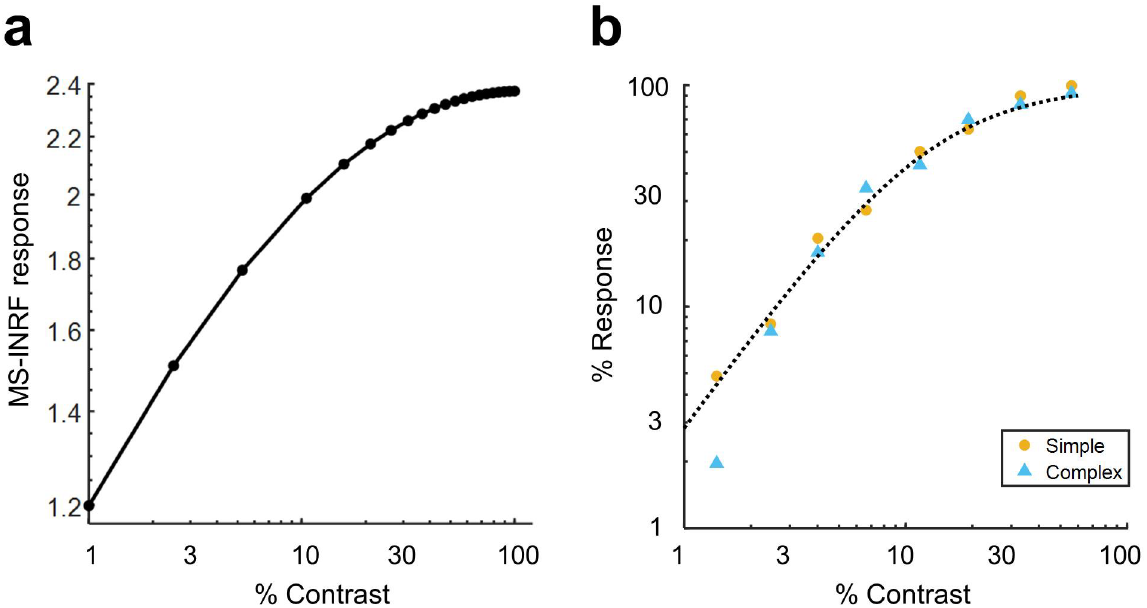
The MS-INRF model reproduces classical findings about contrast saturation. **a**, The magnitude of the MS-INRF response to a sinusoidal grating, as a function of contrast, is monotonically increasing but saturates for high contrast values, as reported in [48, 47, 49]. The response of the MS-INRF model is plotted as a function of stimulus contrast in a log-log scale. Responses are averaged across different presentations with varying spatial phase of the stimuli. **b**, For comparison with **a**, the neural responses of simple and complex cat striate cells to similar stimuli are shown in a log-log scale. Adapted from [49], their figure 11.

#### Motion masking

The proposed model reproduces classical motion masking phenomena [50], where the detection of a moving stimulus is impaired when the signal is masked by a jittering noise that has a similar spatial frequency, see Fig. 5.

**Figure 5:**
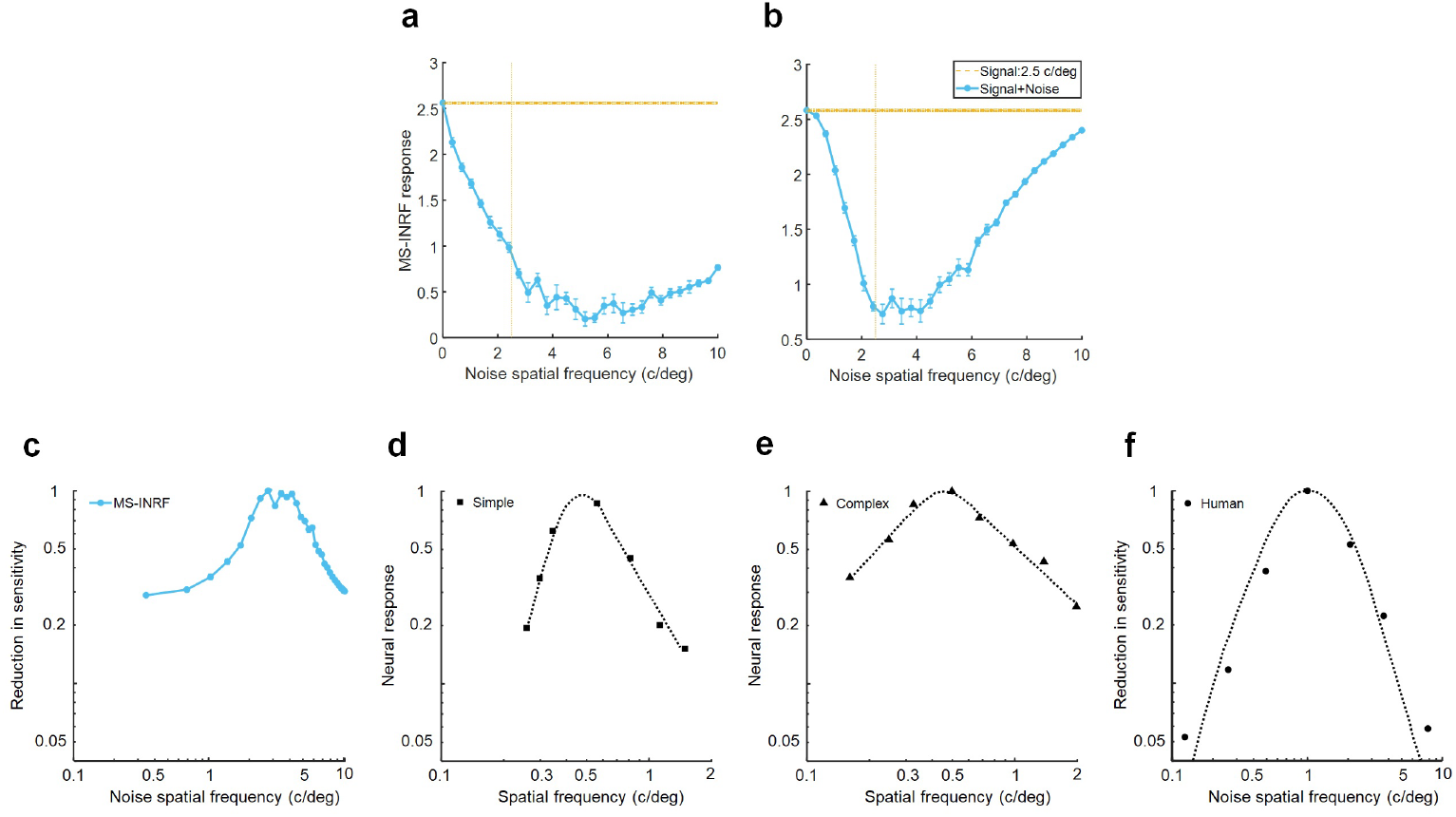
The MS-INRF model reproduces classical findings about motion masking. **a & b**, The MS-INRF response to a signal consisting of a moving sinewave grating (orange) is greatly reduced when we add to the signal jittering noise that has a similar spatial frequency (response to signal plus noise in blue); responses are averaged across different presentations with varying spatial phase of the stimuli. **a**, Simulation of retinal result, where the MS-INRF model is applied directly to the stimulus. **b**, Simulation of cortical result, where the stimulus is filtered with a Difference of Gaussians (DoG) kernel in order to emulate processing in the lateral geniculate nucleus (LGN), and then passed to the MS-INRF model. **c**, Reciprocal of the data shown in **b** (normalized), which may be interpreted as a reduction in sensitivity of the MS-INRF response. The narrow tuning of the masking effect to the spatial frequency of the noise is in agreement with neurophysiological results of simple and complex cell responses (**d & e**, adapted from [70], their figures 3 and 4 respectively) and with psychophysical results observed in humans (**f**, adapted from [50], their figure 1). The phenomenon where the tuning to spatial frequency is narrowed as we add processing stages is consistent with classical physiological results, as retinal ganglion cells are tuned to a broad range of spatial frequencies, LGN cells to a narrower range, and simple V1 cells to an even narrower range of spatial frequencies [70].

#### Reverse-phi motion

If we have a random pattern moving in one direction, but the contrast polarity of the pattern is inverted in every frame, then the perceived direction of motion is the opposite of the actual direction of motion of the pattern. This is a classical phenomenon known as reverse-phi motion [51], and our model reproduces it, as Fig. 6 b shows.

**Figure 6:**
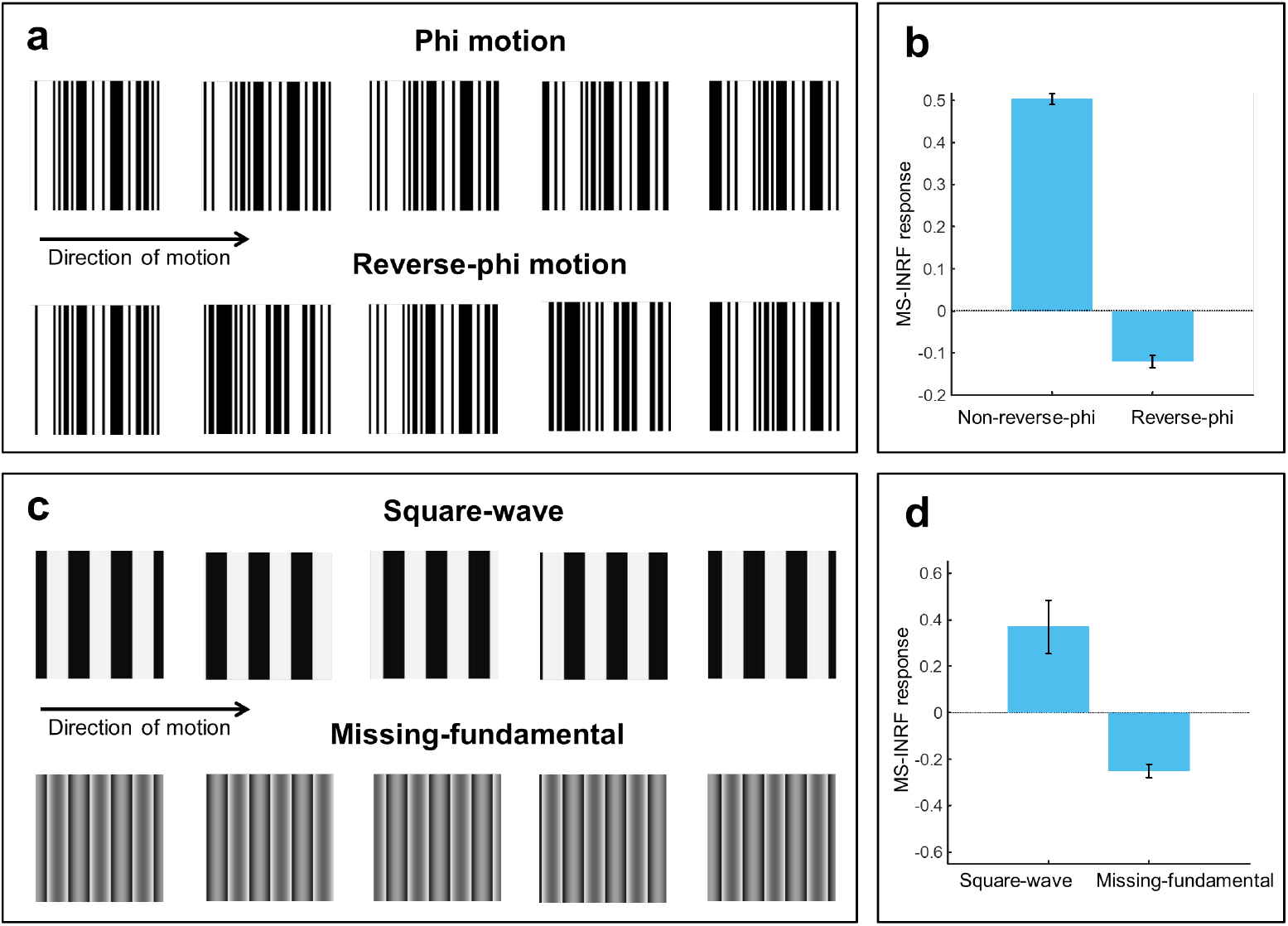
The MS-INRF model reproduces the reverse-phi and the missing fundamental illusions. **a & b**, If we have a random pattern moving in one direction, but its contrast polarity is inverted in every frame, then the perceived direction of motion is the opposite of the actual one. This phenomenon is known as reverse-phi motion [51]. **a**, Top: five frames of a sequence with a random pattern moving rightward (*Phi motion*). Bottom: the same sequence, with rightward motion, but now the contrast polarity is inverted in each frame (*Reverse-phi motion*). **b**, The response of the MS-INRF model to the phi motion stimulus is positive (left bar), indicating rightward motion, while the response of the MS-INRF model to the reverse-phi stimulus is negative (right bar), indicating leftward motion, which shows that the MS-INRF reproduces the perceived inversion of motion of the reverse-phi phenomenon. Model responses are averaged across different random generations of the stimuli. **c & d**, If we have a signal consisting of a square-wave grating moving in a given direction, and we remove its first harmonic from the Fourier series expansion, then the signal is perceived to be moving in the opposite direction: this phenomenon is known as the missing-fundamental illusion [7]. **c**, Top: five frames of a sequence with a square-wave grating moving rightward in quarter-cycle jumps of a certain duration (*Square-wave*). Bottom: the same sequence, with rightward motion, but now the fundamental component (i.e. the first harmonic) has been removed (*Missing-fundamental*). **d**, The response of the MS-INRF model to the square-wave stimulus is positive (left bar), indicating rightward motion, while the response of the MS-INRF model to the missing-fundamental stimulus is negative (right bar), indicating leftward motion, which shows that the MS-INRF reproduces the perceived motion reversal of the missing-fundamental illusion. Responses are averaged across different presentations with varying spatial phase of the stimuli. Also note that if the stimulus moves smoothly instead of in quarter-cycle jumps then the illusion is lost and both signals (square-wave and missing-fundamental) are perceived to move in the same direction, and the MS-INRF model is also reproduces this phenomenon (see Fig. S6 in the supplementary material).

#### Missing-fundamental illusion

If we have a signal consisting of a square-wave grating moving in a given direction, and we remove its first harmonic from the Fourier series expansion, then the signal is perceived to be moving in the opposite direction. This is known as the missing-fundamental illusion [7], and our proposed model is able to reproduce it, as shown in Fig. 6 d.

### 2.3 Properties that are specific to the proposed model

#### The proposed model also detects second-order motion

In Fig. 3 we saw that the MS-INRF model detects first-order motion, and Fig. 7 shows how our model also detects second-order motion. As mentioned above, regular motion models that detect first-order motion are different from models that detect second-order motion, or, put another way, existing motion models for first-order motion are not capable of detecting as well second-order motion without the addenda of some extra processing. Our proposed model, on the other hand, detects both first and second-order motion while keeping the exact same form, i.e. it does not need to change its structure nor its parameters when dealing with one type of motion or the other.

**Figure 7:**
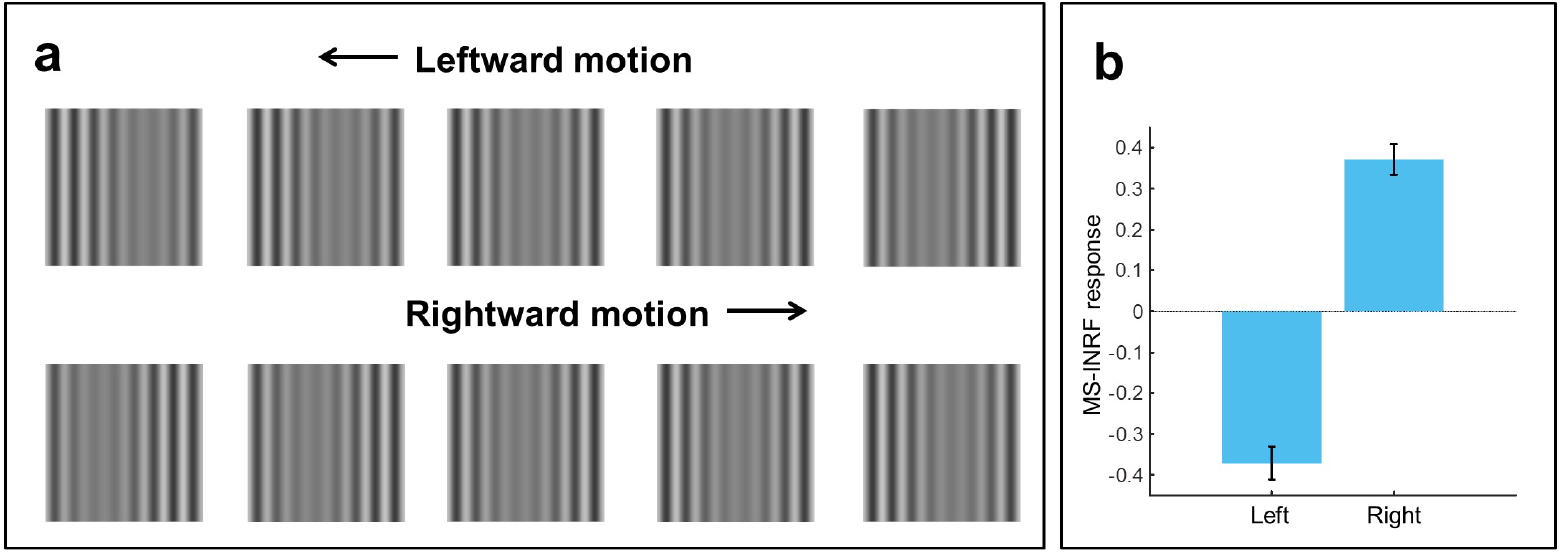
The MS-INRF model is able to detect second-order motion. **a**, Top: five frames of a contrast-defined moving stimulus whose envelope drifts leftward. Bottom: five frames of a contrast-defined moving stimulus whose envelope drifts rightward. **b**, The response of the MS-INRF model to the leftward moving stimulus is negative (left bar), indicating leftward motion, while the response of the MS-INRF model to the rightward moving stimulus is positive (right bar), indicating rightward motion. Responses are averaged across different presentations with varying spatial phase of the stimuli.

#### The proposed model is highly nonlinear

The input/output function of a neuron is often characterized by computing its linear spatio-temporal RF, which, under the assumptions of linear systems theory, represents the stimulus preferred by the neuron, the one to which it is “tuned” [52]. In the same way, in regular motion sensor models, the combination of linear filters and nonlinearities results in a selectivity in space and time. In that context, linear filter theory is used to characterize a motion sensor model by presenting it with sinusoidal gratings covering a wide range of spatial and temporal frequencies. This is a useful representation of the model because the linear systems framework allows us to disregard inputs whose spatio-temporal frequencies fall outside the ranges to which the model is tuned, as they will always produce small or negligible outputs. However, this approach does not make sense for our proposed model due to its highly nonlinear nature, as Fig. 8 shows: there is no spatio-temporal tuning to speak of, because a stimulus of a given spatial and temporal frequency may produce almost no response when presented in isolation but it may affect greatly the output of the model to another stimulus when presented jointly with it (thus violating the linearity assumption).

**Figure 8:**
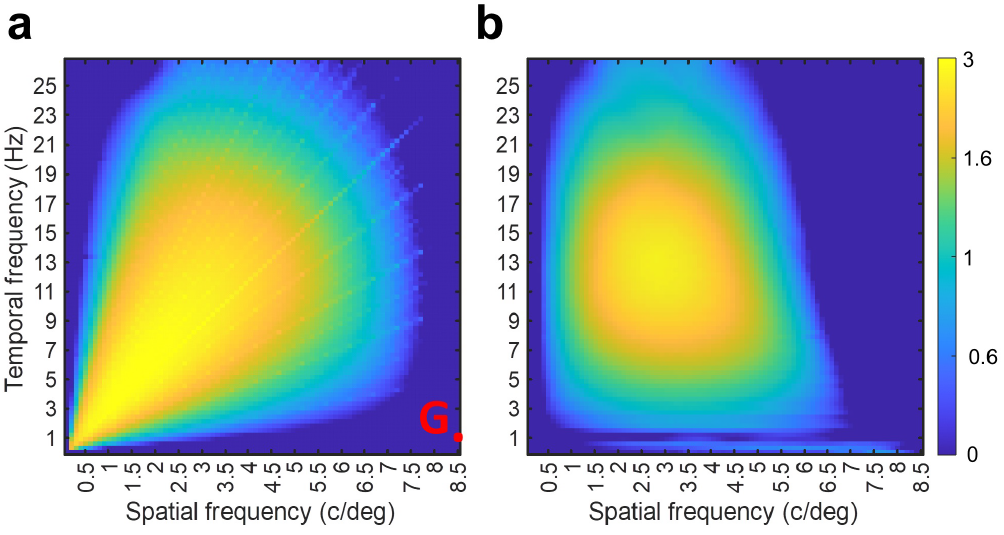
The MS-INRF model is highly nonlinear. **a**, Map of MS-INRF responses to a luminance defined sinusoidal grating as a function of its spatial and temporal frequencies; each point in the map represents a grating, and the point marked with a red dot denotes a grating that we will call *G*, for which the MS-INRF model produces a negligible output. **b**, Map of model responses to stimuli consisting of the sum of grating *G* plus another grating whose spatial and temporal frequencies vary; given that the model response to *G*, when presented in isolation, was negligible, if the model were linear then the map on the right should be identical to the map on the left.

## 3 Discussion

The MS-INRF model incorporates dendritic computations and allows the dendritic nonlinearities to change with the input, which has been shown to be crucial for motion detection for many different species [53, 20] but was not considered by previous motion sensor models. This is not surprising, given that the paramount role of dendrites in shaping neural output was only established after a number of seminal works of the early 2000s [54, 55, 56], and that the contribution of dendrites to sensory perception was confirmed just ten years ago [57].

The highly nonlinear nature of the MS-INRF model is a very desirable property inherited from the INRF formulation, specifically from the input-dependent form of the nonlinear function *σ*, which made that spatial RF model much more powerful in representing nonlinear functions than the standard model (that has a linear RF). This property also allowed the INRF model to remain constant and make accurate predictions for very different inputs, while with the standard model we would need to change the parameters or the model itself [36]. Our results show that the flexibility of the INRF approach also extends to the proposed MS-INRF model, as it can explain data for which classical motion sensor models must resort to the addition of extra stages that increase the model’s complexity and might not be biologically realistic. For instance, the Energy Model explains first-order motion but not the saturation of neural responses with increasing stimulus contrast, a limitation that is solved with the inclusion of additional nonlinear stages [58, 47, 59], and the same happens for detecting second-order motion, for which the model must be extended by interleaving a nonlinearity in between two linear filtering stages [32] (but this new model no longer detects first-order motion). On the other hand our proposed model, just as it comes, works in both scenarios without the need to change it in any way.

The ability of the MS-INRF model to detect both first and second-order motion, and the fact that the proposed motion sensor model is based on a spatio-temporal nonlinear RF model for single neuron activity, have several fundamental implications. Specifically, the MS-INRF model:

- Is a single-stage model, it is not hierarchical; in this way, it overcomes the limitations of current cortical motion sensor models which, as mentioned in the Introduction, are hierarchical, and for this reason they are not consistent with biological evidence showing how motion detection takes place even when intermediate stages in the hierarchy are nulled or bypassed [33, 34].
- Is able to explain second-order motion detection in insects, something that classical algorithms for the retina can not do [15] (also, unlike our model, those algorithms ignore dendritic nonlinearities, when direction selectivity in the retina relies on dendritic computations [21]).
- Is consistent with experimental evidence showing that there are no dedicated second-order motion sensors, while at the same time demonstrating how second-order motion might be detected with the exact same circuitry as for first-order motion, without the need to resort to any additional input (such as motion tracking signals, as proposed in [8]).
- Is consistent with cortical physiology and the sensors it models might in principle be located in any area of the visual cortex, from V1 to MT. This relates the model with the concept of canonical computations in the cortex [35], where neurons located in different cortical areas essentially perform the same type of processing, allowing them in this case to detect both first and second-order motion, as suggested by biological evidence [10, 13].
- Provides a biologically plausible explanation for the surprising algorithmic similarities that insects like the fly and mammals like the rabbit show in motion detection processing [19].

In conclusion, our results suggest that our proposed motion sensor model is very effective as an algorithm, and that its possible neural circuit implementation is biologically compelling at a retinal and cortical level for a variety of species; for these reasons, we believe that our proposed approach is a viable alternative to current models, as it appears to overcome their main limitations.

We want to point out that, whereas the model parameters were manually adjusted so that a single set of values allowed to reproduce all the motion perception phenomena presented in the paper, this manual adjustment process was not intensive and did not require neither exhaustive optimization nor a large number of trials. On the contrary, it was straightforward to identify realistic and suitable parameters for the model, and small variations of the parameter values do not appear to affect the results.

The MS-INRF model can be extended by using different types of dendritic nonlinearities [60], and considering them individually or jointly. We are currently in the process of building a 2D motion model based on combining the signals of several MS-INRF sensors of different preferred orientations, with the aim of explaining challenging 2D motion phenomena such as the perception of plaid motion patterns [61], motion surround suppression [62, 63], direction reversals with the addition of static signals [64], or the sharpening of the speed tuning bandwidth of retinal ganglion cells [65] and MT neurons [66] when exposed to more naturalistic, complex stimulus patterns.

Further work will also include using the stINRF model to develop video quality metrics, for which the INRF approach has shown great promise [67], and replacing linear filters in artificial neural networks with nonlinear stINRF modules, aiming to increase the gains in accuracy, robustness and training data reduction that were already observed with the INRF model [36].

## 4 Methods

### 4.1 MS-INRF model parameters

For all simulations in the paper, we used the same set of parameters for our model (see Eqs. 2 and 3):

The temporal filter *T*_m_ is defined as 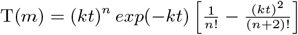, with a total duration of 50 ms, where *t* is the time variable, *k* = 1 and *n* = 5. The temporal filter *T*_*w*_ is defined as 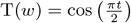, with a total duration of 50 ms. The spatial filter *m* is a Gaussian with a standard deviation of 0.031 deg. The spatial filter *w* is a Gabor oriented vertically, with both the negative and the positive lobe having a horizontal width of 0.125 deg. The weight *λ* is −30. The nonlinearity *σ* is defined as 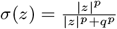, with *p* = 0.4, *q* = 0.1.

### 4.2 Stimuli

All stimuli were generated as sequences of 120 images of 400×400 pixels with values in the range [0,1], where we assume that images span 2 degrees of visual angle and sequences have a duration of 1s (i.e. framerate is 120Hz). Prior to processing by MS-INRF, input stimuli were spatially filtered by the optical blur function proposed in [68]; however, we must note that this step may be omitted without affecting the nature or quality of the results.

Next, we describe the stimuli used to illustrate each of the properties of the MS-INRF formulation.

#### Detection of first-order motion (Fig. 3)

The stimuli consisted of bars of width 0.25 deg moving with a speed of 2 deg/s. See examples of space-time plots for this stimuli in the supplementary material, Fig. S1.

#### Contrast saturation (Fig. 4)

The stimuli consisted of sinusoidal gratings with a spatial frequency of 2 c/deg moving rightward with a temporal frequency of 4 Hz. The stimuli were presented with 20 equally spaced contrasts in the range [0,1] and with 10 equally spaced phases in the range [−*π,π*] (MS-INRF responses were averaged across phases). See examples of space-time plots for this stimuli in the supplementary material, Fig. S2.

#### Motion masking (Fig. 5)

The signal was a sinusoidal grating with Michelson contrast of 0.4 and a spatial frequency of 2.5 c/deg moving rightward with a temporal frequency of 10 Hz; it was presented with 10 equally spaced phases in the range of [−*π,π*], and MS-INRF responses were averaged across phases. The jittering noise was a sinusoidal grating with Michelson contrast of 0.4 and a spatial frequency taking 20 equally spaced values in the range [0,10] c/deg, moving rightward with a temporal frequency of 10 Hz; its phase took random values in the range [−*π,π*] with a frequency of 10 Hz. For the emulation of LGN processing, the stimulus was convolved with a DoG filter where the standard deviation for the center and surround are respectively 0.036 and 0.18 deg, and the balance factor between center and surround is 5 ([69], Eq. (2.45)). See examples of space-time plots for this stimuli (for the isolated signal, the isolated noise, and both combined) in the supplementary material, Fig. S3.

#### Reverse-phi motion (Fig. 6 b)

Both the phi and the reverse-phi stimuli were random stimulus patterns with a Michelson contrast of 0.9, which moved rightward with a speed of 8.5 deg/s. In the case of reverse-phi, contrast polarity was inverted in each frame. MS-INRF responses were averaged across 10 random generations of the stimulus patterns. See examples of space-time plots for this stimuli in the supplementary material, Fig. S4.

#### Missing-fundamental illusion (Fig. 6 d)

Square-wave gratings moved rightward in quarter-cycle jumps that had a duration of 66 ms. Missing-fundamental stimuli were generated by removing from the square-wave gratings the fundamental component (i.e. the first harmonic). Both the square-wave and missing-fundamental stimuli had a Michelson contrast of 0.9, a spatial frequency of 1.5 c/deg, and drifted with a temporal frequency of 4 Hz. They were presented with 10 equally spaced phases in the range [−*π,π*] (MS-INRF responses were averaged across phases). See examples of space-time plots for this stimuli in the supplementary material, Fig. S5.

#### The proposed model also detects second-order motion (Fig. 7)

Contrast-defined gratings were generated drifting rightward and leftward. The envelope had a spatial frequency of 1 c/deg, drifted with a temporal frequency of 7 Hz, and had a Michelson contrast of 0.8. The carrier had a spatial frequency of 4 c/deg and jittered with a temporal frequency of 120 Hz. Modulation depth was 0.3. The modulator was presented with 10 equally spaced phases in the range of [−*π,π*] (MS-INRF responses were averaged across phases). Fig. S7 of the supplementary material shows a space-time plot of a contrast-defined stimulus.

#### The proposed model is highly nonlinear (Fig. 8)

Stimuli integrated by a single sinusoidal component were presented with 100 equally spaced spatial and temporal frequencies in the ranges [0,10] and [0,30] respectively. In the case of compound stimuli, a component was presented with varying spatial and temporal frequencies, exactly as described above, but another component was added that always had a spatial frequency of 8.5 c/deg and a temporal frequency of 1 Hz. Whether compound or integrated by a single component, the gratings moved rightward and each component had a Michelson contrast of 0.4. Components were presented with 10 equally spaced phases in the range of [−*π,π*] (MS-INRF responses were averaged across phases). In the case of compound stimuli, this means a total of 10 *×* 10 = 100 combinations of phases of the two gratings added together. See examples of space-time plots for this stimuli in the supplementary material, Fig. S8.

## Supporting information

Stimulus movies

## 5 Funding

R.L. was supported by a Juan de la Cierva-Formación fellowship (FJC2020-044084-I) funded by Ministerio de Ciencia e Innovación/Agencia Estatal de Investigación (Spain) and by the European Union NextGenerationEU/PRTR. I.S-P was supported by grant PID2021-122245NB-I00, from Ministerio de Ciencia e Innovación (Spain). M.B. was supported by project VIS4NN, Programa Fundamentos 2022, Fundación BBVA, and by grant PID2021-127373NB-I00, Ministerio de Ciencia e Innovacion (Spain).

## 6 Author contributions

All authors have contributed to the conceptualization of the study, development of the software used during simulations, formal analysis and writing.

## 7 Data availability

The MATLAB code implementing the computational motion sensor model (i.e. the MS-INRF model) used to perform the simulations is publicly available at: https://github.com/raullunadelvalle/MS-INRF.

## 8 Competing interests

The authors declare no competing interests.

## Notes

### Competing Interest Statement

The authors have declared no competing interest.

### Summary of Updates

We have changed figures 4 and 5 from the manuscript to incorporate explanations of how the predictions of our computational model match empirical results.

## References

[1] Alexander Borst and Martin Egelhaaf. Principles of visual motion detection. Trends in neurosciences, 12(8):297–306, 1989.

[2] Colin WG Clifford and MR Ibbotson. Fundamental mechanisms of visual motion detection: models, cells and functions. Progress in neurobiology, 68(6):409–437, 2002.

[3] Ken Nakayama. Biological image motion processing: a review. Vision research, 25(5):625–660, 1985.

[4] Jan PH Van Santen and George Sperling. Elaborated reichardt detectors. JOSA A, 2(2):300–321, 1985.

[5] Andrew M Derrington, Harriet A Allen, and Louise S Delicato. Visual mechanisms of motion analysis and motion perception. Annu. Rev. Psychol., 55:181–205, 2004.

[6] Werner Reichardt. Autocorrelation, a principle for evaluation of sensory information by the central nervous system. In Symposium on Principles of Sensory Communication 1959, pages 303–317. MIT press, 1961.

[7] Edward H Adelson and James R Bergen. Spatiotemporal energy models for the perception of motion. Josa a, 2(2):284–299, 1985.

[8] Rémy Allard and Jocelyn Faubert. No dedicated second-order motion system. Journal of Vision, 13(11):2–2, 2013.

[9] Zhong-Lin Lu and George Sperling. Three-systems theory of human visual motion perception: review and update. JOSA A, 18(9):2331–2370, 2001.

[10] Nick Barraclough, Chris Tinsley, Ben Webb, Chris Vincent, and Andrew Derrington. Processing of first-order motion in marmoset visual cortex is influenced by second-order motion. Visual Neuroscience, 23(5):815–824, 2006.

[11] Jutta Billino, Doris I Braun, Frank Bremmer, and Karl R Gegenfurtner. Challenges to normal neural functioning provide insights into separability of motion processing mechanisms. Neuropsychologia, 49(12):3151–3163, 2011.

[12] Howard S Hock and Lee A Gilroy. A common mechanism for the perception of first-order and second-order apparent motion. Vision Research, 45(5):661–675, 2005.

[13] Sang Wook Hong, Frank Tong, and Adriane E Seiffert. Direction-selective patterns of activity in human visual cortex suggest common neural substrates for different types of motion. Neuropsychologia, 50(4):514–521, 2012.

[14] HB Barlow and William R Levick. The mechanism of directionally selective units in rabbit’s retina. The Journal of physiology, 178(3):477, 1965.

[15] Helen H Yang and Thomas R Clandinin. Elementary motion detection in drosophila: algorithms and mechanisms. Annual Review of Vision Science, 4:143–163, 2018.

[16] Jonathan B Demb, Kareem Zaghloul, and Peter Sterling. Cellular basis for the response to second-order motion cues in y retinal ganglion cells. Neuron, 32(4):711–721, 2001.

[17] Alex S Mauss, Anna Vlasits, Alexander Borst, and Marla Feller. Visual circuits for direction selectivity. Annual review of neuroscience, 40:211–230, 2017.

[18] Jamie Carroll Theobald, Brian J Duistermars, Dario L Ringach, and Mark A Frye. Flies see second-order motion. Current Biology, 18(11):R464–R465, 2008.

[19] Damon A Clark and Jonathan B Demb. Parallel computations in insect and mammalian visual motion processing. Current Biology, 26(20):R1062–R1072, 2016.

[20] Susanne E Hausselt, Thomas Euler, Peter B Detwiler, and Winfried Denk. A dendrite-autonomous mechanism for direction selectivity in retinal starburst amacrine cells. PLoS biology, 5(7):e185, 2007.

[21] Alexander Borst, Michael Drews, and Matthias Meier. The neural network behind the eyes of a fly. Current Opinion in Physiology, 16:33–42, 2020.

[22] Yeon Jin Kim, Beth B Peterson, Joanna D Crook, Hannah R Joo, Jiajia Wu, Christian Puller, Farrel R Robinson, Paul D Gamlin, King-Wai Yau, Felix Viana, et al. Origins of direction selectivity in the primate retina. Nature communications, 13(1):2862, 2022.

[23] Thomas D Albright. Form-cue invariant motion processing in primate visual cortex. Science, 255(5048):1141– 1143, 1992.

[24] Daniel J Felleman, Andreas Burkhalter, and David C Van Essen. Cortical connections of areas v3 and vp of macaque monkey extrastriate visual cortex. Journal of Comparative Neurology, 379(1):21–47, 1997.

[25] Luke E Hallum, Michael S Landy, and David J Heeger. Human primary visual cortex (v1) is selective for second-order spatial frequency. Journal of Neurophysiology, 105(5):2121–2131, 2011.

[26] David H Hubel and Torsten N Wiesel. Receptive fields of single neurones in the cat’s striate cortex. The Journal of physiology, 148(3):574–591, 1959.

[27] Guangxing Li, Zhimo Yao, Zhengchun Wang, Nini Yuan, Vargha Talebi, Jiabo Tan, Yongchang Wang, Yifeng Zhou, and Curtis L Baker.Form-cue invariant second-order neuronal responses to contrast modulation in primate area v2. Journal of Neuroscience, 34(36):12081–12092, 2014.

[28] John H Maunsell and David C Van Essen. Functional properties of neurons in middle temporal visual area of the macaque monkey. i. selectivity for stimulus direction, speed, and orientation. Journal of neurophysiology, 49(5):1127–1147, 1983.

[29] Andrew T Smith, Mark W Greenlee, Krish D Singh, Falk M Kraemer, and Jürgen Hennig. The processing of first-and second-order motion in human visual cortex assessed by functional magnetic resonance imaging (fmri). Journal of Neuroscience, 18(10):3816–3830, 1998.

[30] Yi-Xiong Zhou and Curtis L Baker Jr. A processing stream in mammalian visual cortex neurons for non-fourier responses. Science, 261(5117):98–101, 1993.

[31] Robert C Emerson, James R Bergen, and Edward H Adelson. Directionally selective complex cells and the computation of motion energy in cat visual cortex. Vision research, 32(2):203–218, 1992.

[32] Charles Chubb and George Sperling. Two motion perception mechanisms revealed through distance-driven reversal of apparent motion. Proceedings of the National Academy of Sciences, 86(8):2985–2989, 1989.

[33] Pascal Girard, PA Salin, and J Bullier. Response selectivity of neurons in area mt of the macaque monkey during reversible inactivation of area v1. Journal of neurophysiology, 67(6):1437–1446, 1992.

[34] Lawrence C Sincich, Ken F Park, Melville J Wohlgemuth, and Jonathan C Horton. Bypassing v1: a direct geniculate input to area mt. Nature neuroscience, 7(10):1123–1128, 2004.

[35] Kenneth D Miller. Canonical computations of cerebral cortex. Current opinion in neurobiology, 37:75–84, 2016.

[36] Marcelo Bertalmío, Alex Gomez-Villa, Adrián Martín, Javier Vazquez-Corral, David Kane, and Jesús Malo. Evidence for the intrinsically nonlinear nature of receptive fields in vision. Scientific reports, 10(1):1–15, 2020. Nature Publishing Group.

[37] Michael London and Michael Hausser. Dendritic computation. Annu. Rev. Neurosci., 28:503–532, 2005.

[38] R Angus Silver. Neuronal arithmetic. Nature Reviews Neuroscience, 11(7):474–489, 2010.

[39] Yuri Elias Rodrigues, Cezar M Tigaret, Hélène Marie, Cian O’Donnell, and Romain Veltz. A stochastic model of hippocampal synaptic plasticity with geometrical readout of enzyme dynamics. Elife, 12:e80152, 2023.

[40] Ilias Rentzeperis, Dario Prandi, and Marcelo Bertalmío. A neural model for v1 that incorporates dendritic nonlinearities and back-propagating action potentials. bioRxiv, 2024.

[41] Marcelo Bertalmío, Alexia Durán Vizcaíno, Jesús Malo, and Felix A Wichmann. Plaid masking explained with input-dependent dendritic nonlinearities. Scientific Reports, 14(1):24856, 2024.

[42] Bruno A Olshausen. 20 years of learning about vision: Questions answered, questions unanswered, and questions not yet asked. In 20 Years of Computational Neuroscience, pages 243–270. Springer, 2013.

[43] Karl J Friston. The labile brain. iii. transients and spatio–temporal receptive fields. Philosophical Transactions of the Royal Society of London. Series B: Biological Sciences, 355(1394):253–265, 2000.

[44] Y Fukushima, K Hara, and M Kimura. A spatio-temporal model of ganglion cell receptive field in the cat retina. Biological cybernetics, 54(2):91–98, 1986.

[45] Albert Gidon, Timothy Adam Zolnik, Pawel Fidzinski, Felix Bolduan, Athanasia Papoutsi, Panayiota Poirazi, Martin Holtkamp, Imre Vida, and Matthew Evan Larkum. Dendritic action potentials and computation in human layer 2/3 cortical neurons. Science, 367(6473):83–87, 2020.

[46] Ádám Magó, Noémi Kis, Balazs Lükő, and Judit K Makara. Distinct dendritic ca2+ spike forms produce opposing input-output transformations in rat ca3 pyramidal cells. Elife, 10:e74493, 2021.

[47] David J Heeger. Normalization of cell responses in cat striate cortex. Visual neuroscience, 9(2):181–197, 1992.

[48] Izumi Ohzawa, Gary Sclar, and Ralph D Freeman. Contrast gain control in the cat’s visual system. Journal of neurophysiology, 54(3):651–667, 1985.

[49] Duane G Albrecht and David B Hamilton. Striate cortex of monkey and cat: contrast response function. Journal of neurophysiology, 48(1):217–237, 1982.

[50] Stephen J Anderson and David C Burr. Spatial and temporal selectivity of the human motion detection system. Vision research, 25(8):1147–1154, 1985.

[51] SM Anstis. Phi movement as a subtraction process. Vision research, 10(12):1411–IN5, 1970.

[52] Matteo Carandini, Jonathan B Demb, Valerio Mante, David J Tolhurst, Yang Dan, Bruno A Olshausen, Jack L Gallant, and Nicole C Rust. Do we know what the early visual system does? Journal of Neuroscience, 25(46):10577–10597, 2005.

[53] Alexander Borst, Jürgen Haag, and Alex S Mauss. How fly neurons compute the direction of visual motion. Journal of Comparative Physiology A, 206(2):109–124, 2020.

[54] Michael Häusser and Bartlett Mel. Dendrites: bug or feature? Current opinion in neurobiology, 13(3):372–383, 2003.

[55] Panayiota Poirazi, Terrence Brannon, and Bartlett W Mel. Pyramidal neuron as two-layer neural network. Neuron, 37(6):989–999, 2003.

[56] Alon Polsky, Bartlett W Mel, and Jackie Schiller. Computational subunits in thin dendrites of pyramidal cells. Nature neuroscience, 7(6):621, 2004.

[57] Spencer L Smith, Ikuko T Smith, Tiago Branco, and Michael Häusser. Dendritic spikes enhance stimulus selectivity in cortical neurons in vivo. Nature, 503(7474):115–120, 2013.

[58] David J Heeger. Half-squaring in responses of cat striate cells. Visual neuroscience, 9(5):427–443, 1992.

[59] David J Heeger. Modeling simple-cell direction selectivity with normalized, half-squared, linear operators. Journal of neurophysiology, 70(5):1885–1898, 1993.

[60] Panayiota Poirazi and Athanasia Papoutsi. Illuminating dendritic function with computational models. Nature Reviews Neuroscience, 21(6):303–321, 2020.

[61] Edward H Adelson and J Anthony Movshon. Phenomenal coherence of moving visual patterns. Nature, 300(5892):523–525, 1982.

[62] Davis M Glasser and Duje Tadin. Low-level mechanisms do not explain paradoxical motion percepts. Journal of Vision, 10(4):20–20, 2010.

[63] Ignacio Serrano-Pedraza, Ellen L Hogg, and Jenny CA Read. Spatial non-homogeneity of the antagonistic surround in motion perception. Journal of vision, 11(2):3–3, 2011.

[64] Andrew M Derrington and G Bruce Henning. Errors in direction-of-motion discrimination with complex stimuli. Vision Research, 27(1):61–75, 1987.

[65] César R Ravello, Laurent U Perrinet, María-José Escobar and Adrián G Palacios. Speed-selectivity in retinal ganglion cells is sharpened by broad spatial frequency, naturalistic stimuli. Scientific reports, 9(1):456, 2019.

[66] Nicholas J Priebe, Carlos R Cassanello, and Stephen G Lisberger. The neural representation of speed in macaque area mt/v5. Journal of Neuroscience, 23(13):5650–5661, 2003.

[67] Raúl Luna, Itziar Zabaleta, and Marcelo Bertalmío. State-of-the-art image and video quality assessment with a metric based on an intrinsically non-linear neural summation model. Frontiers in Neuroscience, 17, 2023.

[68] JJ Vos, TJTP Van den Berg, et al. Report on disability glare. CIE collection, 135(1):1–9, 1999.

[69] Peter Dayan, Laurence F Abbott, et al. Theoretical neuroscience: computational and mathematical modeling of neural systems. Journal of Cognitive Neuroscience, 15(1):154–155, 2003.

[70] Lamberto Maffei and Adriana Fiorentini. The visual cortex as a spatial frequency analyser. Vision research, 13(7):1255–1267, 1973.

