## Supplementary material for "Overcoming the limitations of motion sensor models by considering dendritic computations"

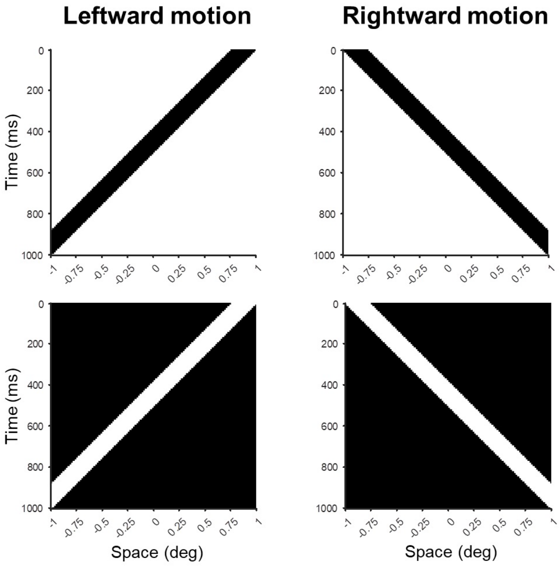


Figure S1: **Space-time plots of moving bars.** The upper panels show black bars moving over a white background and the lower panels show white bars moving over a black background. The left panels show bars moving leftward and right panels show bars moving rightward. The bars have a spatial width of 0.25 deg and move with a speed of 2 deg/s.

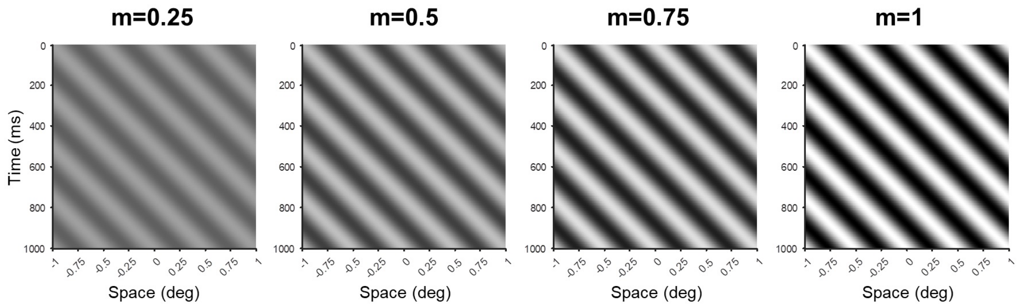


Figure S2: **Space-time plots of luminance-defined gratings with varying Michelson contrast.** Constrast is, from left to right: m=0.25, m=0.5, m=0.75 and m=1. The stimuli have a spatial frequency of 2 c/deg and move rightward with a temporal frequency of 4 Hz. They are depicted with a phase of 0 deg.

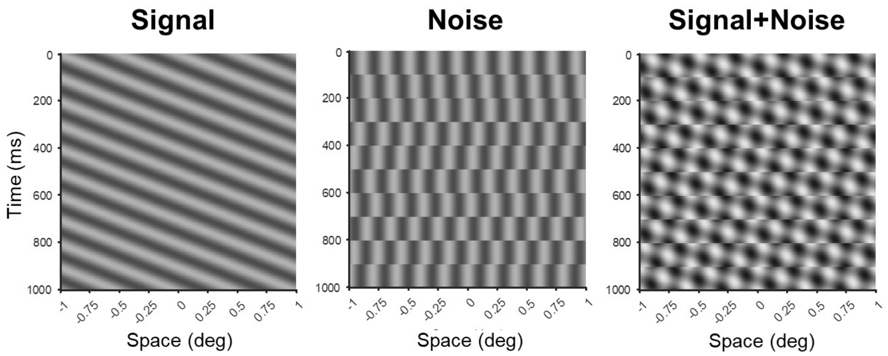


Figure S3: **Space-time plots for signal, a noise and a signal+noise stimuli.** Signal luminance-defined grating (left panel), jittering noise (middle panel) and the addition of both (right panel). The signal has a spatial frequency of 2.5 c/deg, is shown with a phase of 0 deg and drifts rightward with a temporal frequency of 10 Hz. The noise has a spatial frequency of 5 c/deg and jitters with a temporal frequency of 10 Hz too. Both, signal and noise, have a Michelson contrast of 0.4.

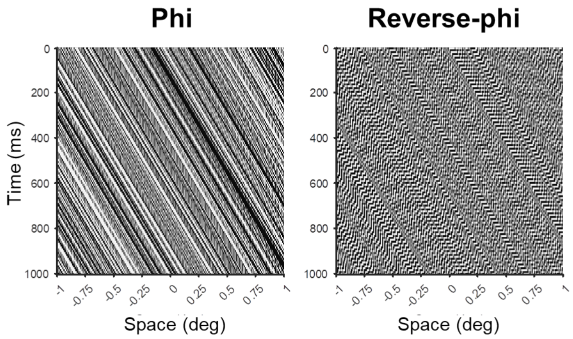


Figure S4: **Space-time plots of a phi and a reverse-phi stimulus.** Luminance-defined noise moving rightward (Phi; left panel) and the resulting pattern when contrast polarity is inverted in each frame (Reverse-phi; right panel). In this example, the stimulus patterns drift with a speed of 1 deg/s, and have a Michelson contrast of 0.9.

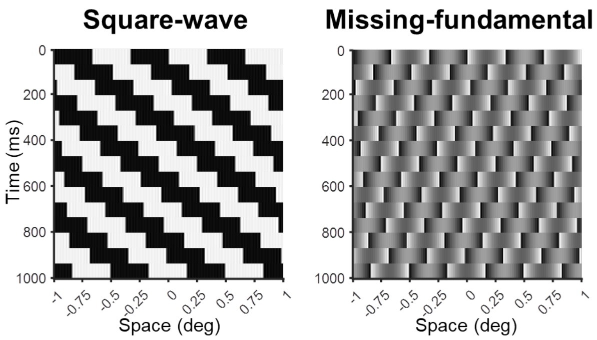


Figure S5: **Space-time plots of a square-wave and a missing-fundamental stimulus**. Square-wave grating moving rightward in quarter-cycle jumps of 66 ms duration (left panel), and that stimulus when its fundamental component (i.e. first harmonic) is removed (right panel). The stimuli have a Michelson contrast of 0.9, a spatial frequency of 1.5 c/deg, and drift with a temporal frequency of 4 Hz.

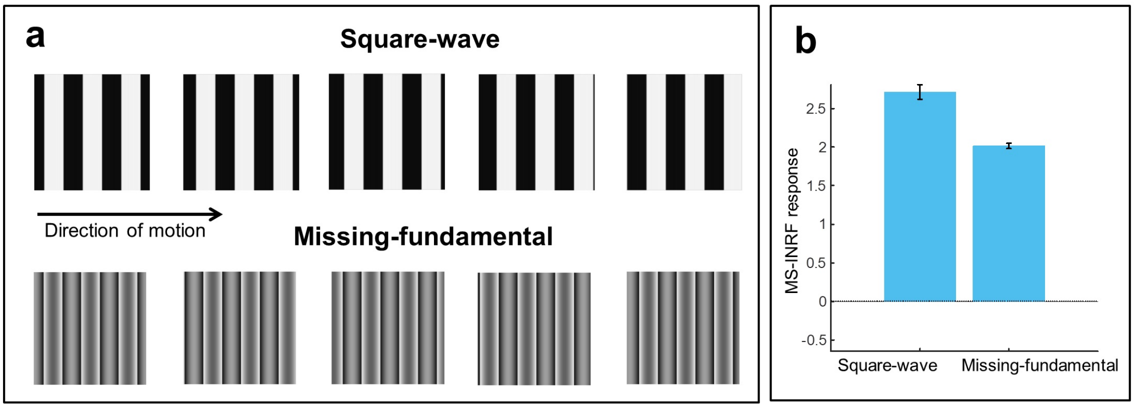


Figure S6: **The MS-INRF model also reproduces the case when the missing-fundamental illusion fails.** If we have a signal consisting of a square-wave grating moving in a given direction, and we remove its first harmonic from the Fourier series expansion, then the signal is perceived to be moving in the opposite direction: this is the missing-fundamental illusion, but it only works if the stimuli moves in jumps, otherwise the illusion is lost and both signals (square-wave and missing-fundamental) are perceived to move in the same direction. **a**, Top: five frames of a sequence with a square-wave grating smoothly moving to the right (*Square-wave*). Bottom: the same sequence, with rightward motion, but now the fundamental component (i.e. the first harmonic) has been removed from the signal (*Missing-fundamental*). **b**, The response of the MS-INRF model to the square-wave stimulus is positive (left bar), indicating rightward motion, and the response of the MS-INRF model to the missing-fundamental stimulus is also positive (right bar), which shows that the MS-INRF reproduces the loss of the missing-fundamental illusion. Responses are averaged across different presentations with varying spatial phase of the stimuli.

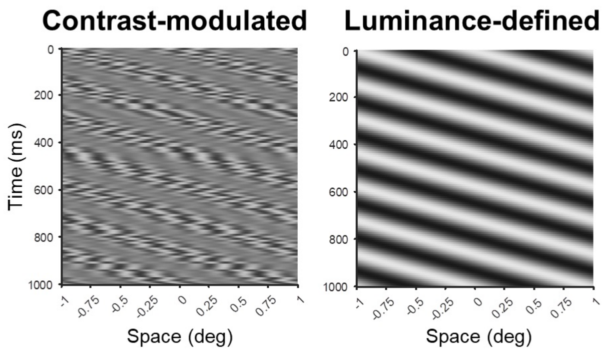


Figure S7: **Space-time plots of a contrast-modulated and a luminance-defined stimulus.** Contrast-modulated stimulus (left panel), where the envelope has a spatial frequency of 1 c/deg, drifts rightward with 7 Hz temporal frequency, and has a Michelson contrast of 0.8. It is depicted with a phase of 0 deg. The carrier has a spatial frequency of 4 c/deg and jitters with a temporal frequency of 120 Hz. Modulation depth is 0.3. The right panel shows the space-time plot of a luminance-defined grating for comparison. As well, it has a spatial frequency of 1 c/deg, drifts rightward with 7 Hz temporal frequency, has a Michelson contrast of 0.8 and is depicted with a phase of 0 deg.

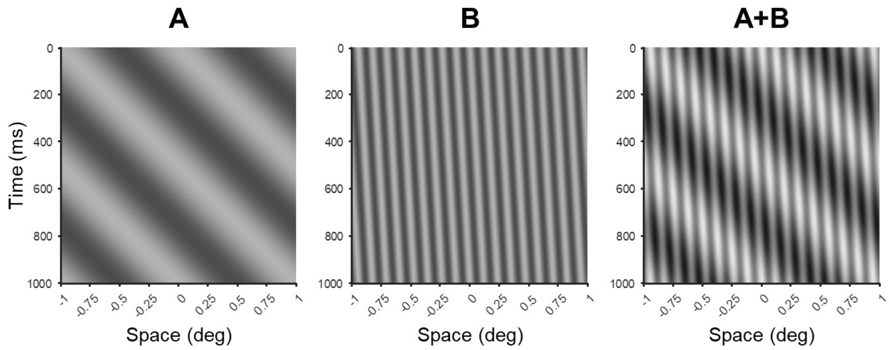


Figure S8: **Space-time plots of simple luminance-defined gratings and the compound stimulus resulting from their addition.** Luminance-defined grating of 1 c/deg spatial frequency drifting rightward with a temporal frequency of 2 Hz (A; left panel); luminance-defined grating of 8.5 c/deg spatial frequency drifting rightward with a temporal frequency of 1 Hz (A; middle panel); and the compound stimulus resulting from adding the two previous components (A+B; right panel). The components have a Michelson contrast of 0.4 and are depicted with a phase of 0 deg.

### Stimulus movies

The file "Stimulus_movies" has example movies with the only purpose of better understanding the stimuli used in this work, and for this reason some of their parameters may not match exactly those of the Figures to which they relate. Below, we associate each movie file with the Figure it exemplifies:

- Fig1a_First_Order: First-order motion stimulus in Fig. 1a. of the main text.
- Fig1b_Second_Order: Second-order motion stimulus in Fig. 1b. of the main text.
- FigS1_LeftwardMotion_BlackOverWhite: Leftward-moving black bar over a white background in Fig. S1.
- FigS1_LeftwardMotion_WhiteOverBlack: Leftward-moving white bar over a black background in Fig. S1.
- FigS1_RightwardMotion_BlackOverWhite: Rightward-moving black bar over a white background in Fig. S1.

- FigS1_RightwardMotion_WhiteOverBlack: Rightward-moving white bar over a black background in Fig. S1.
- FigS2_m_0p25: Luminance-defined grating with 0.25 Michelson contrast in Fig. S2.
- FigS2_m_0p5: Luminance-defined grating with 0.5 Michelson contrast in Fig. S2.
- FigS2_m_0p75: Luminance-defined grating with 0.75 Michelson contrast in Fig. S2.
- FigS2_m_1: Luminance-defined grating with 1 Michelson contrast in Fig. S2.
- FigS3_Signal: Signal stimulus in Fig. S3.
- FigS3_Noise: Noise stimulus in Fig. S3.
- FigS3_SignalPlusNoise: Signal+Noise stimulus in Fig. S3.
- FigS4_Phi: Phi stimulus in Fig. S4.
- FigS4_Reverse-phi: Reverse-phi stimulus in Fig. S4.
- FigS5_Square-wave: Square-wave grating in Fig. S5.
- FigS5_Missing-fundamental: Missing-fundamental stimulus in Fig. S5.
- FigS7_Contrast-modulated: Contrast-modulated stimulus in Fig. S7.
- FigS7_Luminance-defined: Luminance-defined stimulus in Fig. S7.
- FigS8_A: Luminance-defined grating A in Fig. S8.
- FigS8_B: Luminance-defined grating B in Fig. S8.
- FigS8_APlusB: Luminance-defined grating made by adding together A and B in Fig. S8.
